# Methionine Alkylation as an Approach to Quantify Methionine Oxidation Using Mass Spectrometry

**DOI:** 10.1101/2023.09.21.558873

**Authors:** Margaret Hoare, Ruiyue Tan, Kevin A. Welle, Kyle Swovick, Jennifer R. Hryhorenko, Sina Ghaemmaghami

## Abstract

Post-translational oxidation of methionine residues can destabilize proteins or modify their functions. Although levels of methionine oxidation can provide important information regarding the structural integrity and regulation of proteins, their quantitation is often challenging as analytical procedures in and of themselves can artifactually oxidize methionines. Here, we develop a mass spectrometry-based method called Methionine Oxidation by Blocking with Alkylation (MObBa) that quantifies methionine oxidation by selectively alkylating and blocking unoxidized methionines. Thus, alkylated methionines can be used as a stable proxy for unoxidized methionines. Using proof of concept experiments, we demonstrate that MObBa can be used to measure methionine oxidation levels within individual synthetic peptides and on proteome-wide scales. MObBa may provide a straightforward experimental strategy for mass spectrometric quantitation of methionine oxidation.

**Significance Statement:** Over time, cellular proteins can become oxidatively damaged by reactive oxygen species (ROS). A residue that is particularly prone to oxidative damage is methionine. Here, we develop and validate a methodology for detecting and quantifying levels of methionine oxidation by mass spectrometry. This approach has a number of practical advantages over methods currently available for analysis of methionine oxidation. The ability to accurately quantify methionine oxidation will provide important insights into factors that influence protein homeostasis within a cell.

## Introduction

Methionine is a sulfur-containing amino acid that is susceptible to oxidation.^1, 2^ Sidechain oxidation converts nonpolar methionine residues to polar methionine sulfoxides, and this change in hydrophobicity can dramatically alter the structure and function of proteins.^3–7^ The conversion of methionine to methionine sulfoxide can occur chemically through reactions with reactive oxygen species (ROS), or enzymatically through the action of specific oxygenases.^8–11^ Observed levels of methionine oxidation *in vivo* are also influenced by cellular activities of a class of reducing enzymes known as methionine sulfoxide reductases (Msrs).^12^ As a post-translational modification, methionine oxidation has been implicated in protein damage, cellular signaling, ROS scavenging, pathological aging, and etiology of several neurodegenerative diseases.^6, 13–15^

Techniques for quantifying methionine oxidation include FT-IR spectroscopy, chromatography and mass spectrometry.^16–23^ Among these, methods involving tandem mass spectrometry (LC-MS/MS) are particularly advantageous as they enable measurements of methionine oxidation on proteome-wide scales.^17, 18, 22, 24^ For example, a protocol referred to as COFRADIC takes advantage of changes in liquid chromatography (LC) peptide retention times in bottom-up proteomic experiments to globally quantify methionine sulfoxide levels.^22, 24^ However, a general complication with mass spectrometric analysis of methionine oxidation is that spontaneous oxidation during sample preparation and ionization steps of proteomic workflows often results in unpredictable background accumulation of methionine sulfoxides.^25, 26^ Hence, because of methionines’ propensity for spurious oxidation, it is often difficult to unequivocally differentiate between methionine sulfoxides that form *in vivo* and those that artifactually accumulate during sample preparation and subsequent mass spectrometric analyses.^18^

An alternative mass spectrometric approach that circumvents the artifactual oxidation of methionines is Methionine Oxidation by Blocking (MObB).^16–18, 21^ In MObB, unoxidized methionines within peptides are forcibly oxidized with excess levels of ^18^O-labeled hydrogen peroxide, thus preventing further spontaneous ^16^O oxidation during mass spectrometric analysis. Accurate oxidation levels can subsequently be determined by measuring relative ratios of ^18^O– to ^16^O-modified peptide levels following a bottom-up proteomic workflow. Although MObB provides a straightforward proteomic approach for quantitation of methionine oxidation, it has a number of experimental complications. First, because ^18^O– and ^16^O-labeled peptides vary in mass by only 2 Daltons, their spectral isotopic envelopes are overlapped and measuring their relative levels requires non-standard data analysis procedures.^17, 18^ Second, the quantitative accuracy of the method is contingent on the high isotopic purity of ^18^O-labeled hydrogen peroxide. Third, ^18^O– labeled hydrogen peroxide is a costly reagent that can be difficult to obtain. In fact, at the time of writing this manuscript, the previously available commercial sources for ^18^O-H_2_O_2_ (Sigma and Cambridge Isotope Laboratories) had discontinued its sale. Thus, it would be advantageous to develop an alternative method that can selectively block and quantify methionines using more standard and commonly available reagents. Here, we describe a protocol named Methionine Oxidation by Blocking with Alkylation (MObBa) for selective alkylation and quantitation of unoxidized methionines by iodoacetamide (IAA). In comparison to ^18^O-H_2_O_2,_ IAA is significantly less expensive, commonly available, and amenable to standard proteomic workflows and data analysis. We determine the precision of this technique and assess its ability to quantify methionine oxidation levels on proteome-wide scales.

## Results

### Methionine can be fully alkylated by iodoacetamide at low pH

Although the alkylation of cysteine thiols is a common practice in proteomic workflows, the alkylation of methionines has been comparatively less explored.^27–30^ It has long been known that methionines can be selectively alkylated at a pH of 2-5 with iodoacetate (IA) and iodoacetamide (IAA).^31–35^ More recently, conjugation of methionines with a range of other modifying reagents has also been described.^36, 37^ The selective alkylation of unoxidized methionines, as detected by shifts in native gel electrophoretic mobility, has previously been used as an approach to quantify levels of protein oxidation in intact proteins.^31, 35^ Here, we take advantage of the fact that methionines (but not methionine sulfoxides) can be alkylated at low pH to develop a method for quantitation of methionine oxidation in bottom-up proteomic experiments.

In MObBa, polypeptides are treated with IAA at low pH to specifically modify unoxidized methionines and block further oxidation (Figure 1A). Fractional oxidation levels are then determined by measuring peptide alkylation levels relative to fully reduced Msr-treated controls. We initially set out to validate the chemical premise of MObBa using a methionine-containing synthetic peptide. Specifically, we verified that 1) methionines can be fully alkylated at low pH with IAA, 2) methionine oxidation inhibits its alkylation, and 3) methionine alkylation inhibits its oxidation. We carried out alkylation reactions at pH 4 over the course of 7 days on a synthetic methionine-containing peptide (MASILKKLAVDR) (see methods for details). The efficiency of alkylation was determined by mass spectrometry. The data indicate that methionine can be fully alkylated by IAA and this alkylation is completely inhibited by oxidation (Figure 1B). The kinetics of this reaction at 37 °C is relatively slow and full alkylation is achieved after 4 days. The alkylation of the peptide completely inhibits its subsequent oxidation by H_2_O_2_, indicating that it is an effective blocking strategy (Figure 1C).

**Figure 1.**
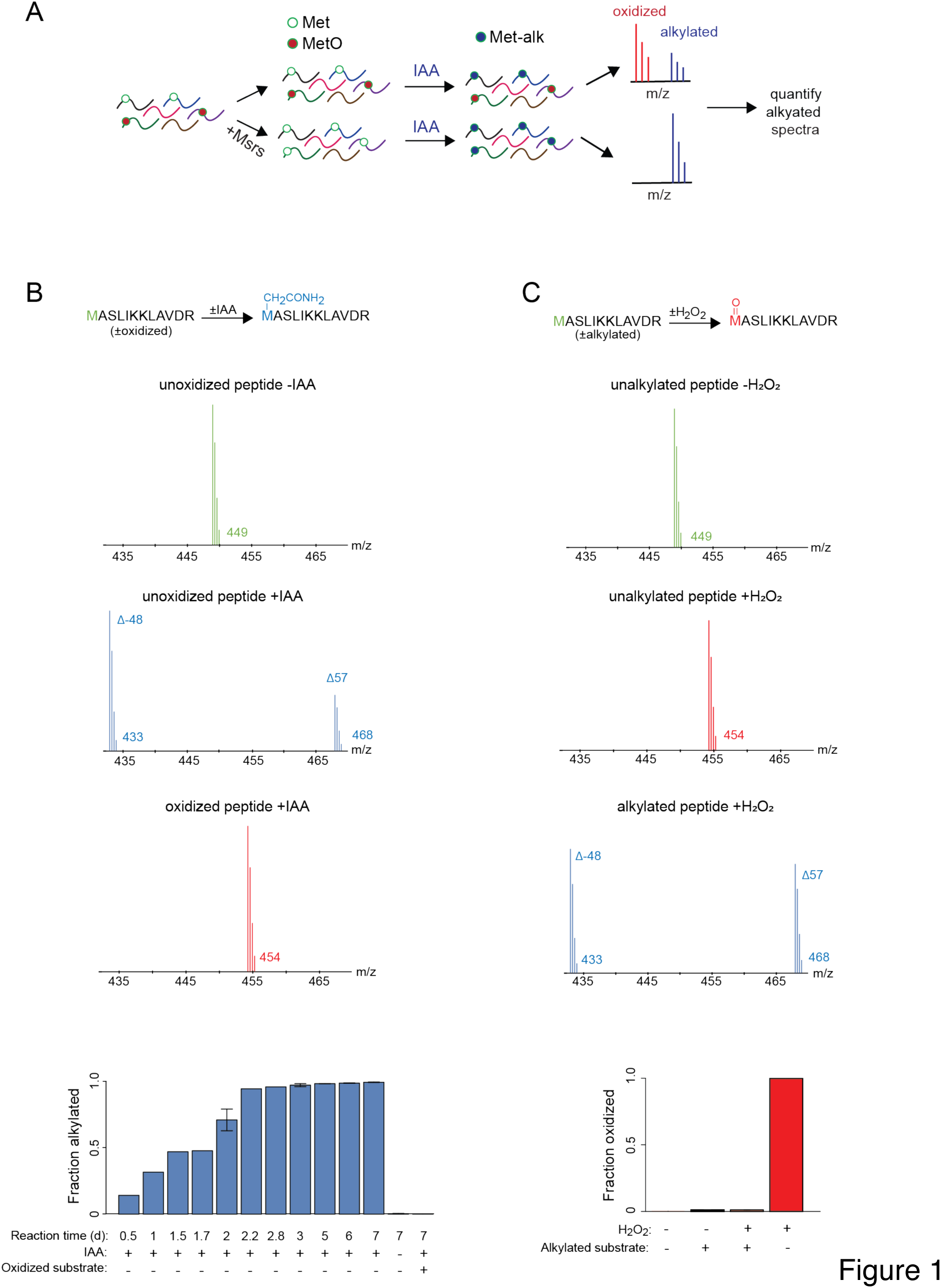
Unoxidized methionines can be fully alkylated by iodoacetamide. **A)** The overall workflow of MObBa. Unoxidized methionines are selectively alkylated by iodoacetamide (IAA). This alkylation prevents further spurious oxidation and can be quantified by mass spectrometry as a proxy the for fraction of methionines that are unoxidized in the sample. **B)** The synthetic peptide MASLIKKLAVDR can be fully alkylated by IAA and this alkylation is inhibited by methionine oxidation. Oxidized or unoxidized peptides were used as substrates as indicated. For the experiments that generated the three illustrated spectra, peptides were either treated with IAA at pH 4 for 7 days, or left untreated. The green, blue and red spectra, indicate the expected m/z of the +3 charge state of the unoxidized, alkylated (including both Δ57 and Δ-48 modifications), and oxidized forms of the peptide. The bar plot shows the time-course of the alkylation reaction. Fractional alkylation was quantified by adding the total intensities of the carbamidomethylation (Δ57) and dethiomethylation (Δ-48) modifications and dividing by the total intensity of the peptide. The error bars indicate the standard deviations of 2 replicate experiments. **C)** The alkylation of the synthetic peptide MASLIKKLAVDR inhibits subsequent methionine oxidation. Alkylated or unalkylated peptides were used as substrates as indicated. For the experiments that generated the three illustrated spectra, peptides were either treated with 160 mM H_2_O_2_ at pH 5 for 30 minutes, or left untreated. The spectra are colored according to the scheme described in B. The bar plot shows the quantitation of fractional oxidation as measured by dividing the intensity of the oxidized peptide by the total intensity. The error bars indicate the standard deviations of 2 replicate experiments.

It had been previously reported that the alkylation of methionines by IAA yields two detectable modified peptide products in LC-MS/MS experiments.^28, 30^ The first is a +57 Da product generated by carbamidomethylation of the sidechain (Δ57), and the second is a –48 Da neutral loss product resulting from the alkylation-induced dethiomethylation of the sidechain (Δ-48). Both of these products were detectable in the alkylated samples and together accounted for the entire population of modified peptides (Figure 1B,C).

### Methionine alkylation by iodoacetamide is selective

We next determined whether methionine can be selectively alkylated without artifactual modification of non-sulfur containing residues. A fully unoxidized peptide mixture was generated from an *E. coli* protein extract by reducing disulfide bonds, alkylating free cystines with IAA at neutral pH, and digesting the sample with trypsin. The sample was then treated with IAA at low pH for 7 days. As cysteines were already fully alkylated, they were not further modified by this treatment. Treated samples were analyzed by LC–MS/MS and searched against the *E. coli* proteome. Levels of modification for each amino acid were determined by quantifying the number of peptides harboring the unmodified form of each residue (Figure 2A). The data indicated that IAA almost fully depleted peptides containing unmodified methionines, while relative levels of peptides containing unmodified forms of other residues were largely unaffected. In a second experiment, the peptide mixture was initially reduced by treatment with Msrs and then treated with IAA as above. LC-MS/MS analysis indicated that nearly all identified methionine-containing peptides in IAA-treated samples harbored either methionine carbamidomethylation (Δ57) or dethiomethylation (Δ-48) (Figure 2B).

**Figure 2.**
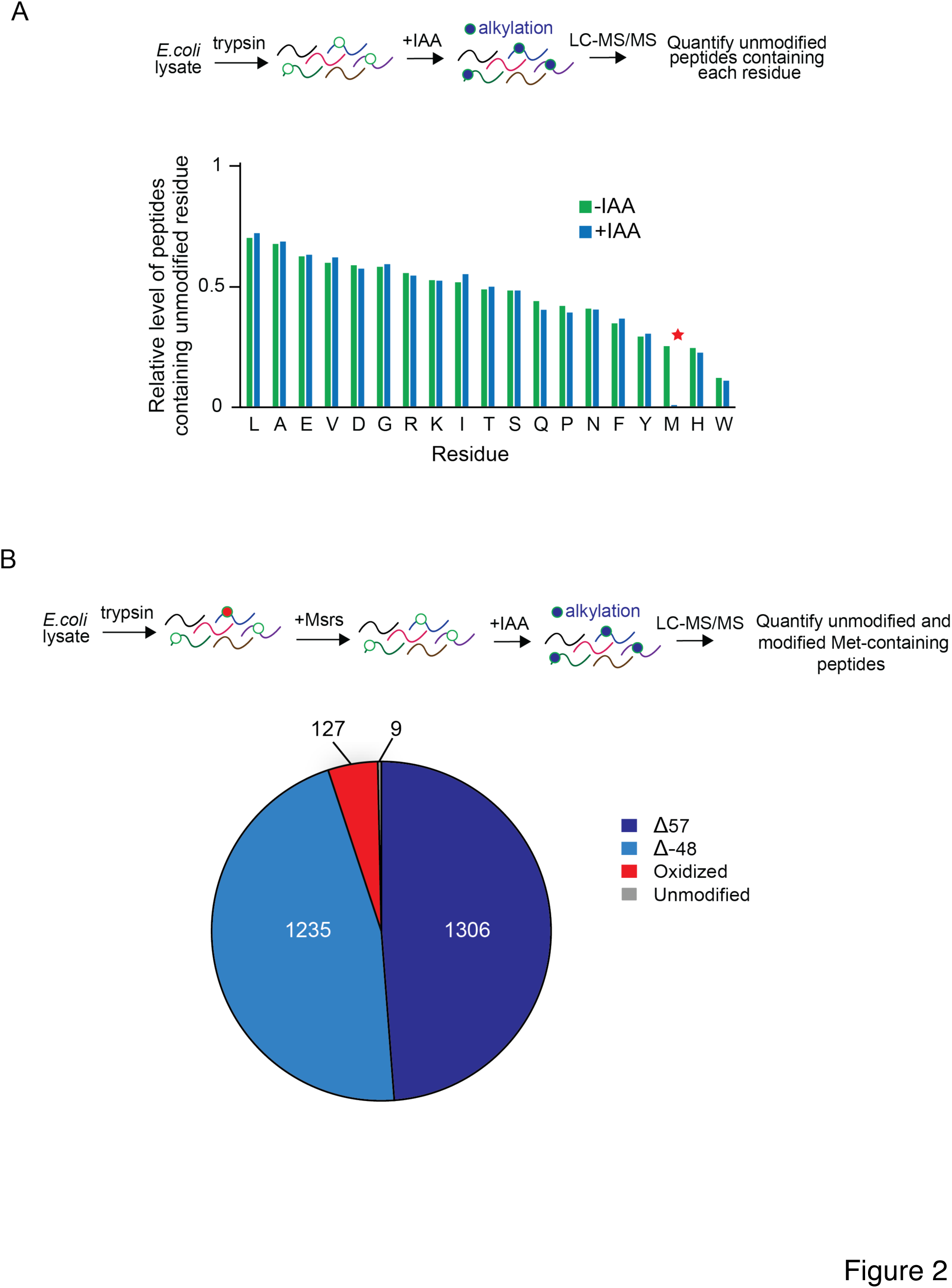
Methionine can be selectively alkylated by iodoacetamide in proteome-wide experiments. **A)** Peptide mixtures generated by trypsinization of *E. coli* extracts were left untreated or incubated with IAA at pH 4 and analyzed by LC-MS/MS. Numbers of unique peptides containing each residue (in their unmodified forms) were divided by the number of all detected peptides to determine the relative population of each unmodified residue within the peptide population. Cysteines were not included as they are pre-alkylated prior to the low pH IAA treatment. **B)** Tryptic peptides generated from *E. coli* extracts were treated with Msrs to remove background methionine oxidation then incubated with IAA and analyzed by LC-MS/MS to determine the number of unique methionine-containing peptides in their unmodified, dethiomethylated (Δ-48), carbamidomethylated (Δ57) and oxidized forms.

### MObBa accurately measures protein oxidation levels

To demonstrate the quantitative accuracy of MObBa, we generated and analyzed peptide mixtures with known levels of methionine oxidation. A methionine-containing synthetic peptide was fully oxidized by the addition of H_2_O_2_, or completely reduced by addition of Msrs (Figure 3). The two samples were mixed at variable ratios to obtain different known levels of methionine oxidation. The mixtures were then treated with IAA and oxidation levels were quantified by summing the intensities of Δ57 and Δ-48 modified peptides. The results indicated a high level of quantitative accuracy and a linear response across the range of analyzed fractional oxidation levels.

**Figure 3.**
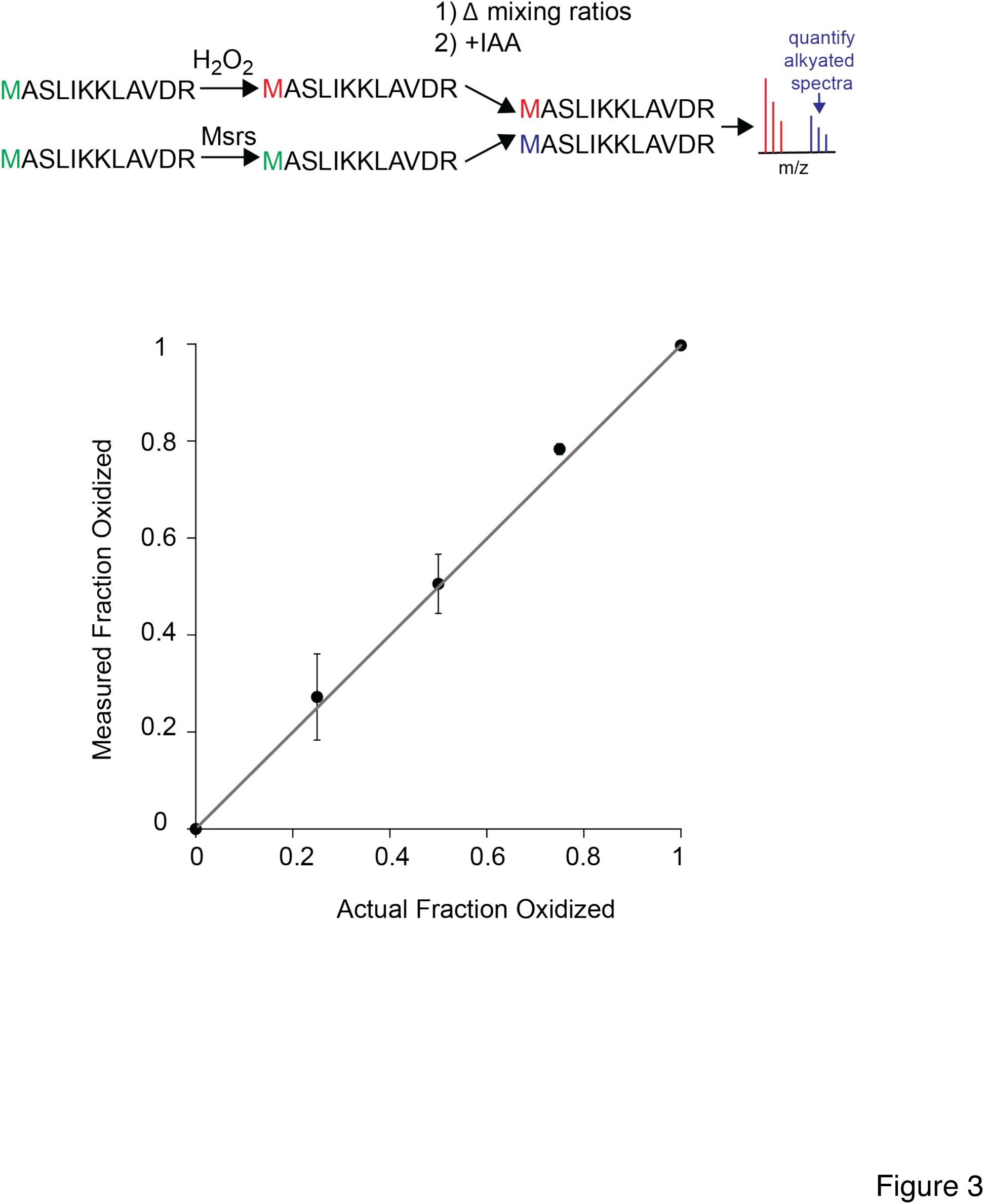
MObBa can accurately quantify methionine oxidation levels for a synthetic peptide. The synthetic peptide MASLIKKLAVDR was exposed to two treatments: full oxidation of methionine with hydrogen peroxide and full reduction of methionine oxidation by Msrs. Mixtures of the two treatments (containing 0%, 25%, 50%, 75%, 100% oxidized peptides) were incubated with IAA and analyzed by mass spectrometry. For each mixture, fractional oxidation was measured by calculating the relative intensities of unalkylated peptides after IAA treatment. The error bars indicate the standard deviations of 2 replicate experiments.

We next repeated the above experiment on the entire *E. coli* proteome (Figure 4). Proteins were extracted from *E. coli* and digested into peptides by trypsin after reduction and blocking of cysteines. The peptide mixture was divided into two different treatment conditions: 1) full reduction of oxidized methionines by Msrs and 2) full oxidation of methionines by hydrogen peroxide. Peptides from these two conditions were mixed to generate specific fractional oxidation levels. Each mixture was then treated with IAA at low pH, analyzed by LC-MS/MS, and searched against the *E. coli* proteome using the Δ57 and Δ-48 alkylated forms as variable modifications. Fractional oxidation levels were measured by normalizing the intensities of alkylated peptides relative to unoxidized controls. The results indicated that the distribution and median oxidation measurements of peptides corresponded to expected levels within each mixture.

**Figure 4.**
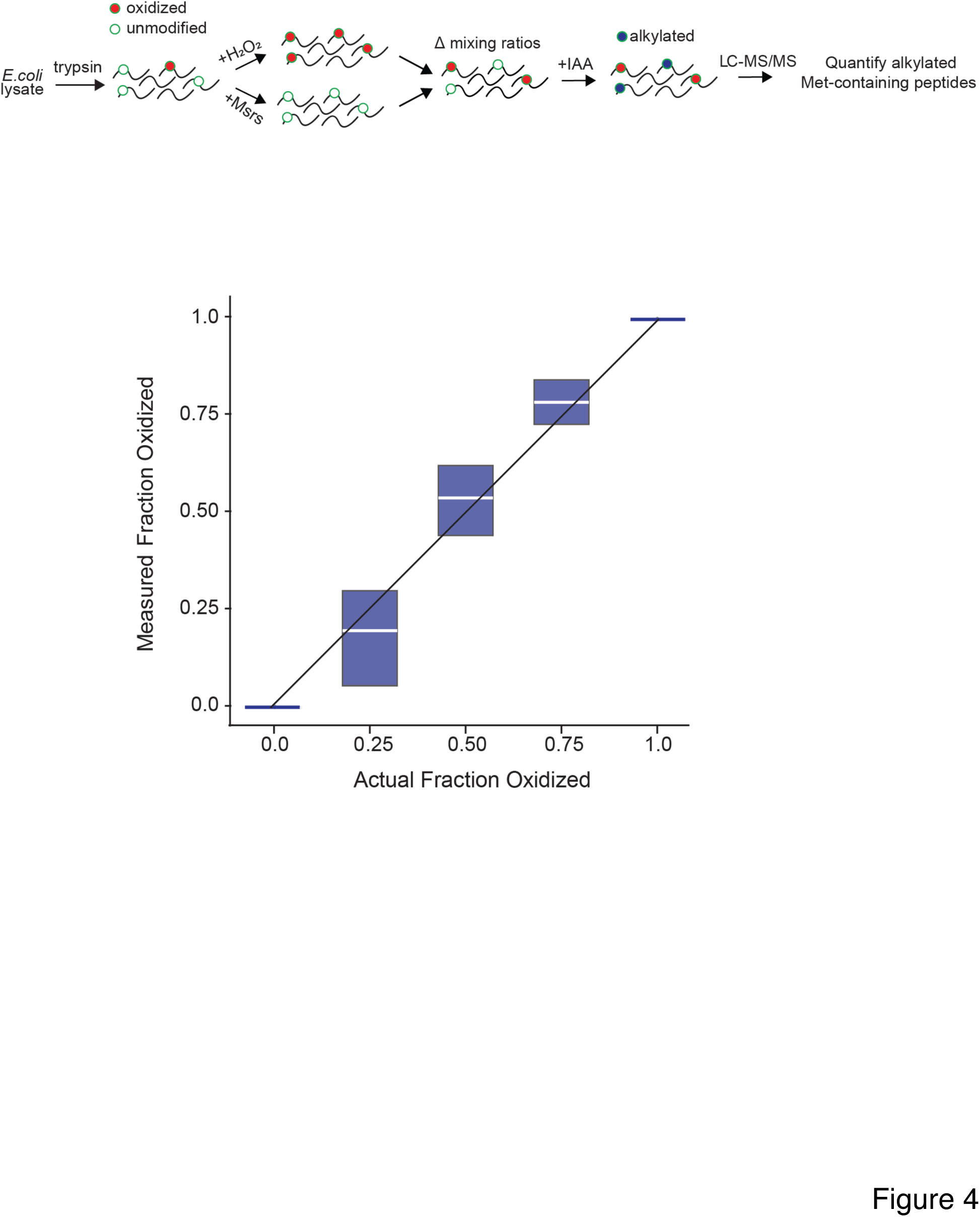
MObBa accurately measures methionine oxidation in proteome-wide experiments. Tryptic peptides derived from *E. coli* were exposed to two treatments: full oxidation by hydrogen peroxide and full reduction by Msrs. Mixtures of the two treatments (containing 0%, 25%, 50%, 75%, and 100% oxidized peptides) were incubated with IAA and analyzed by LC-MS/MS. Fractional oxidation of each detected peptide was measured as in Figure 3. The box plot indicates the interquartile distribution of the measurements in each condition and the white line indicates the median.

## Discussion

An important consideration for accurate quantitation of methionine oxidation by mass spectrometry is the prevention of artifactual oxidation that occurs during the sample preparation and ionization steps of typical proteomics workflows. In this study, we show that treatment of peptides with IAA at low pH alkylates unoxidized methionines and prevents their subsequent oxidation during mass spectrometric analysis. Thus, quantitation of alkylated methionines following IAA treatment provides an effective metric for levels of oxidized methionines that were present prior to alkylation.

The experiments presented here validate MObBa as an approach for quantifying methionine oxidation in both isolated polypeptides and complex peptide mixtures. They also highlight two important caveats that must be considered when implementing this strategy. First, the alkylation of methionines is a relatively slow reaction, occurring over the course of 7 days at 37 °C. The long reaction time not only adds to the overall analysis time but may also lead to accumulation of oxidation prior to alkylation. In the experiments described above, we minimized this effect through careful degassing of all samples and by conducting reactions under nitrogen gas. The second potential complication of MObBa is that peptide ions harboring carbamidomethylated methionines appear to be unstable in electrospray mass spectrometry and result in a neutral loss product where the methionine sidechain is dethiomethylated. Thus, to account for all blocked peptide products, both carbamidomethylated and dethiomethylated modified forms of methionine must be included in database searches. The sidechain neutral loss also prohibits the possible use of isotopically labeled IAA as an approach to compare alkylation levels across samples.^38^ However, as an alternative, it may be possible to combine MObBa with metabolic labeling (e.g. SILAC) to measure differences in oxidation levels between peptide samples obtained from cells exposed to different treatment conditions.

Despite the above-mentioned limitations, MObBa offers a number of advantages compared to alternative methods for quantitation of methionine oxidation. It is a straightforward protocol involving IAA, a commonly used reagent that is part of the typical bottom-up proteomics workflow. Analysis of data generated by MObBa does not require specialized software and can be conducted using commonly available search algorithms. Thus, MObBa may prove useful for analyses that require quantitation of methionine oxidation without requiring unconventional procedures and reagents.

## Materials and methods

### Peptide preparation

The synthetic peptide MASLIKKLAVDR was purchased from GenScript at 97.7% purity. The *E. coli* peptide extract was prepared from an *E. coli* K12 W3110 strain. Cells were grown in LB media at 37 °C, pelleted, then lysed in 5% SDS in 50 mM TEAB through sonication at 25 amps. Cellular debris was pelleted by centrifugation at 16,000 xg for 15 minutes. Protein concentrations were quantified from the supernatant using a BCA kit (Thermo Scientific). 25 µg of protein was reduced with 2 mM dithiothreitol (Sigma) for 60 minutes at 55 °C. Cysteines were alkylated with 10mM iodoacetamide (Sigma) for 30 minutes at room temperature in the dark. The alkylation reaction was quenched with 1.2% phosphoric acid. 90% methanol in 100 mM TEAB was added to the extract (6:1,v/v) and the sample was loaded onto S-trap micro filters (ProtiFi). The peptides were isolated in the filters and then digested with 1 µg of trypsin (Pierce) and incubated overnight at 37°C in a water bath. After incubation, the filters were centrifuged for 1 minute at 4000 xg, then eluted with 40 µl of 0.1% trifluoroacetic acid (TFA) in H_2_O (Thermo Scientific) and then 40 µl of 50% ACN/H_2_O in 0.1% TFA (Thermo Scientific). The peptides were lyophilized and reconstituted in degassed 5% formic acid (FA) in H_2_O.

### Alkylation of peptide-bound methionines

For the synthetic peptide, 15 µg of peptide was incubated with 50 µl of 33 mM iodoacetamide (Sigma) in degassed 5% formic acid (pH 4) under nitrogen gas at 37 °C. Unless otherwise stated, the reaction was carried out for 7 days. The samples were then desalted in a homemade C18 column and eluted in 50% ACN/H_2_O in 0.1% formic acid. *E. coli* peptide extracts were alkylated as above after reconstitution in 5% FA.

### Oxidation of peptide-bound methionines

For both synthetic peptides and *E.coli* extracts, 25 µg of peptide was oxidized with 50 µl of 160 mM H_2_O_2_ for 30 minutes at 37 °C. The sample was then frozen and lyophilized to remove excess hydrogen peroxide, then desalted in a homemade C18 column and eluted in 50% ACN/H_2_O in 0.1% formic acid.

### Expression and purification of MsrA and MsrB

pET 151/ D-TOPO expression vectors coding for *E. coli* Methionine Sulfoxide Reductase A and B containing N-terminal His tags downstream of a T7 promoter were synthesized (Invitrogen GeneArt) and transformed into BL21(DE3) competent cells (Thermo Scientific) by heat shock. Cells were plated and placed under ampicillin selection overnight at 37°C. A 10 mL culture of LB media was inoculated with single colony and grown to an OD_600_ of 0.6-0.8. The culture was subsequently added to 1 L of LB media and induced with 400 µM IPTG, then incubated overnight at 25 °C while shaking at 180 rpm. The bacteria were pelleted by centrifugation at 8,000 xg for 3 minutes. The cells were placed in PBS buffer containing 20 mM imidazole and EDTA free protease inhibitor at pH 7.4 and sonicated. The lysate was centrifuged at 6,000 xg for 30 minutes at 4°C and the pellet was discarded. A nickel column was made with Ni-NTA resin (Thermo Scientific) and equilibrated with two washes of 20 mM sodium phosphate and 10 mM imidazole in PBS at pH 7.4. The lysate was run through the column by gravity flow. The column was then washed with 25 mM imidazole at pH 7.4 until no protein was detected in the flowthrough by UV absorption. Bound proteins were eluted with 250mM imidazole in pH 7.4. Dialysis was performed to transfer proteins into degassed 50mM Tris buffer at pH 7.4. Protein concentrations were determined by bicinchoninic acid (BCA) assay (Thermo Scientific). The final purity of the MsrA and MsrB enzymes were ∼91% and ∼94%, respectively, as determined by SDS-PAGE. The activity of the purified enzymes was verified by measuring reduction of oxidized peptides as detected by mass spectrometry (see below).

### Reduction of peptide-bound methionine sulfoxides by Msrs

For the synthetic peptide, 25 µg of the peptide was incubated with 50 mM dithiothreitol (Sigma), 1.5 µg of MsrA and 5.0 µg of MsrB in 50mM Tris buffer at pH 7.4 for 45 minutes at 37 °C. Samples were lyophilized then desalted to remove enzymes and salts. *E. coli* peptide extracts were reduced as above.

### Mass spectrometric analysis

Synthetic peptides were diluted to 15 µg/mL in 50% ACN/H_2_O in 0.1% FA. 50 µl of the sample was run in a Q Exactive Plus Mass Spectrometer (Thermo Scientific) by direct injection with a Dionex Ultimate 3000 with a flow rate of 100 µL/min for 3 minutes. The solvent consisted of a 50% mixture of 0.1% FA in H_2_O and 0.1% FA in ACN. Peptides were ionized by a HESI source set in positive mode. Data were collected over a range of 300-2000 m/z at a resolution of 70K at m/z 200 with a 240 ms maximum injection time and AGC target of 1e6.

*E. coli* peptide extracts were injected onto a homemade 30cm C18 column with 1.8 uM beads (Sepax), with an Easy nLC-1200 HPLC (Thermo Fisher), connected to a Fusion Lumos Tribrid mass spectrometer (Thermo Fisher).

Solvent A was 0.1% formic acid in water while solvent B was 0.1% formic acid in 80% acetonitrile. Ions were introduced to the mass spectrometer using a Nanospray Flex source operating at 2 kV. The gradient began at 3% B and held for 2 minutes, increased to 10% B over 5 minutes, increased to 38% B over 68 minutes, then ramped up to 90% over 3 minutes and held for 3 minutes, before returning to starting conditions over 2 minutes and re-equilibrating the column for 7 minutes, for a total run time of 90 minutes. For all experiments, the Fusion Lumos was operated in data-dependent mode with Advanced Peak Determination (ADP) set to “TRUE” and Monoisotopic Precursor Selection (MIPS) set to “Peptide”. The full MS1 scan was done over a range of 375-1400 m/z with a resolution of 120K at m/z of 200, an AGC target of 4e5, and a maximum injection time of 50 ms. Peptides with a charge state between 2-5 were chosen for fragmentation. Dynamic exclusion was set to 20 seconds and to exclude after 1 time with low and high mass tolerances of 10 ppm. For the experiments shown in Figure 2A, MS1 and MS2 scans were acquired in the Orbitrap (OT) and ion trap (IT) respectively with a cycle time of 1.5 seconds. Precursor ions were fragmented by collision induced dissociation (CID) using a collision energy of 30%, an activation time of 10 ms, an activation Q of 0.25, and with an isolation window of 1.1 m/z. The IT scan rate was set to “Rapid” with a maximum ion injection time of 35 ms and an AGC target of 1e4. The minimum and maximum intensity thresholds were set to 2e4 and 1e20 respectively. For the experiments shown in Figures 2B and 4, MS1 and MS2 scans were performed in the OT with a cycle time of 2 seconds. Precursor ions were fragmented by higher energy collision dissociation (HCD) using a collision energy of 30% and isolation width of 1.1 m/z. MS2 scans were performed at 15K resolution at m/z of 200 with an AGC target of 5e4 and maximum injection time of 25 ms.

### Measurement of fractional oxidation

For synthetic peptide experiments, raw MS data were analyzed by the XCalibur software (Thermo Scientific). The total intensity of the peptides containing the alkylation modifications were summed and fractional alkylation was calculated in the experimental sample. For figures 1 and 3, MS1 spectra were exported using the MSConvert software and intensities of alkylated or oxidized peaks were measured using Mathematica (Wolfram).

Proteome-wide data obtained by the Fusion Lumos samples were searched in Proteome Discoverer (Thermo Fisher) against the *E. coli* reference proteome downloaded in April 2019. Precursor mass and fragment mass tolerances were set to 10 ppm and 0.6 Da respectively. Cysteine carbamidomethylation (+57.021 Da) was a fixed modification and carbamidomethylation (+57.021 Da), dethiomethylation (−48.003 Da), oxidation (+15.995 Da), and methionine-loss (−131.040 Da) were variable modifications for methionine. The oxidation level analyses were conducted using the measured PSM intensities as described in the Results section.

Raw spectra and search results for all experiments have been deposited in the PRIDE database (Acc: PXD045497).

## Author Information

### Corresponding Author

*

### Author Contributions

The study concept was conceived by M.H., R.T. and S.G. The experiments were mostly carried out by M.H. Mass spectrometry was performed by K.W., K.S. and J.H. Data analysis was conducted by M.H., K.W. and S.G. The initial draft of the manuscript was written by M.H. and S.G.

## Data Availability

All raw and processed data are available in the included Supporting Information and at the ProteomeXchange Consortium via the PRIDE partner repository (accession number PXD045497). Currently, the data can be accessed with the username and password xaIpHm9s.

## Abbreviations

MObBa: Methionine Oxidation by Blocking with Alkylation
ROS: reactive oxygen species
Msrs: methionine sulfoxide reductases
MObB: Methionine Oxidation by Blocking
LC-MS/MS: liquid chromatography tandem mass spectrometry
LC: liquid chromatography
IAA: iodoacetamide
IA: iodoacetate
Δ57: methionine carbamidomethylation
Δ-48: methionine dethiomethylation
SILAC: Stable Isotopic Labeling by Amino Acids in Cell Culture
DDA: data-dependent acquisition
MIPS: Monoisotopic Precursor Selection
CID: collision-induced dissociation

## References

1. Kim, G.; Weiss, S. J.; Levine, R. L., Methionine oxidation and reduction in proteins. Biochim Biophys Acta 2014, 1840 (2), 901–5.

2. Walker, E. J.; Bettinger, J. Q.; Welle, K. A.; Hryhorenko, J. R.; Molina Vargas, A. M.; O’Connell, M. R.; Ghaemmaghami, S., Protein folding stabilities are a major determinant of oxidation rates for buried methionine residues. J Biol Chem 2022, 298 (5), 101872.

3. Chao, C. C.; Ma, Y. S.; Stadtman, E. R., Modification of protein surface hydrophobicity and methionine oxidation by oxidative systems. Proc Natl Acad Sci U S A 1997, 94 (7), 2969–74.

4. Hsu, Y. R.; Narhi, L. O.; Spahr, C.; Langley, K. E.; Lu, H. S., In vitro methionine oxidation of Escherichia coli-derived human stem cell factor: effects on the molecular structure, biological activity, and dimerization. Protein Sci 1996, 5 (6), 1165–73.

5. Samson, A. L.; Knaupp, A. S.; Kass, I.; Kleifeld, O.; Marijanovic, E. M.; Hughes, V. A.; Lupton, C. J.; Buckle, A. M.; Bottomley, S. P.; Medcalf, R. L., Oxidation of an exposed methionine instigates the aggregation of glyceraldehyde-3-phosphate dehydrogenase. J Biol Chem 2014, 289 (39), 26922–26936.

6. Bettinger, J.; Ghaemmaghami, S., Methionine oxidation within the prion protein. Prion 2020, 14 (1), 193–205.

7. Nakaso, K.; Tajima, N.; Ito, S.; Teraoka, M.; Yamashita, A.; Horikoshi, Y.; Kikuchi, D.; Mochida, S.; Nakashima, K.; Matsura, T., Dopamine-mediated oxidation of methionine 127 in alpha-synuclein causes cytotoxicity and oligomerization of alpha-synuclein. PLoS One 2013, 8 (2), e55068.

8. Manta, B.; Gladyshev, V. N., Regulated methionine oxidation by monooxygenases. Free Radic Biol Med 2017, 109, 141–155.

9. Hung, R.-J.; Pak, C. W.; Terman, J. R., Direct Redox Regulation of F-Actin Assembly and Disassembly by Mical. Science 2011, 334 (6063), 1710–1713.

10. Grintsevich, E. E.; Ge, P.; Sawaya, M. R.; Yesilyurt, H. G.; Terman, J. R.; Zhou, Z. H.; Reisler, E., Catastrophic disassembly of actin filaments via Mical-mediated oxidation. Nature Communications 2017, 8 (1), 2183.

11. Terman, J. R.; Mao, T.; Pasterkamp, R. J.; Yu, H. H.; Kolodkin, A. L., MICALs, a family of conserved flavoprotein oxidoreductases, function in plexin-mediated axonal repulsion. Cell 2002, 109 (7), 887–900.

12. Moskovitz, J., Methionine sulfoxide reductases: ubiquitous enzymes involved in antioxidant defense, protein regulation, and prevention of aging-associated diseases. Biochim Biophys Acta 2005, 1703 (2), 213–9.

13. Levine, R. L.; Moskovitz, J.; Stadtman, E. R., Oxidation of methionine in proteins: roles in antioxidant defense and cellular regulation. IUBMB Life 2000, 50 (4-5), 301–7.

14. Stadtman, E. R.; Van Remmen, H.; Richardson, A.; Wehr, N. B.; Levine, R. L., Methionine oxidation and aging. Biochim Biophys Acta 2005, 1703 (2), 135–40.

15. Lockhart, C.; Smith, A. K.; Klimov, D. K., Methionine Oxidation Changes the Mechanism of Abeta Peptide Binding to the DMPC Bilayer. Sci Rep 2019, 9 (1), 5947.

16. Liu, H.; Ponniah, G.; Neill, A.; Patel, R.; Andrien, B., Accurate Determination of Protein Methionine Oxidation by Stable Isotope Labeling and LC-MS Analysis. Analytical Chemistry 2013, 85 (24), 11705–11709.

17. Bettinger, J. Q.; Simon, M.; Korotkov, A.; Welle, K. A.; Hryhorenko, J. R.; Seluanov, A.; Gorbunova, V.; Ghaemmaghami, S., Accurate Proteomewide Measurement of Methionine Oxidation in Aging Mouse Brains. Journal of Proteome Research 2022, 21 (6), 1495–1509.

18. Bettinger, J. Q.; Welle, K. A.; Hryhorenko, J. R.; Ghaemmaghami, S., Quantitative Analysis of in Vivo Methionine Oxidation of the Human Proteome. Journal of Proteome Research 2020, 19 (2), 624–633.

19. Balakrishnan, G.; Barnett, G. V.; Kar, S. R.; Das, T. K., Detection and Identification of the Vibrational Markers for the Quantification of Methionine Oxidation in Therapeutic Proteins. Analytical Chemistry 2018, 90 (11), 6959–6966.

20. Houde, D.; Kauppinen, P.; Mhatre, R.; Lyubarskaya, Y., Determination of protein oxidation by mass spectrometry and method transfer to quality control. Journal of Chromatography A 2006, 1123 (2), 189–198.

21. Shipman, J. T.; Go, E. P.; Desaire, H., Method for Quantifying Oxidized Methionines and Application to HIV-1 Env. Journal of the American Society for Mass Spectrometry 2018, 29 (10), 2041–2047.

22. Ghesquière, B.; Gevaert, K., Proteomics methods to study methionine oxidation. Mass Spectrometry Reviews 2014, 33 (2), 147–156.

23. Lee, B. C.; Péterfi, Z.; Hoffmann, F. W.; Moore, R. E.; Kaya, A.; Avanesov, A.; Tarrago, L.; Zhou, Y.; Weerapana, E.; Fomenko, D. E.; Hoffmann, P. R.; Gladyshev, V. N., MsrB1 and MICALs Regulate Actin Assembly and Macrophage Function via Reversible Stereoselective Methionine Oxidation. Molecular Cell 2013, 51 (3), 397–404.

24. Ghesquière, B.; Jonckheere, V.; Colaert, N.; Van Durme, J.; Timmerman, E.; Goethals, M.; Schymkowitz, J.; Rousseau, F.; Vandekerckhove, J.; Gevaert, K., Redox Proteomics of Protein-bound Methionine Oxidation. Molecular & Cellular Proteomics: MCP 2011, 10 (5), M110.006866.

25. Zang, L.; Carlage, T.; Murphy, D.; Frenkel, R.; Bryngelson, P.; Madsen, M.; Lyubarskaya, Y., Residual metals cause variability in methionine oxidation measurements in protein pharmaceuticals using LC-UV/MS peptide mapping. J Chromatogr B Analyt Technol Biomed Life Sci 2012, 895-896, 71–6.

26. Chen, M.; Cook, K. D., Oxidation artifacts in the electrospray mass spectrometry of Abeta Peptide. Anal Chem 2007, 79 (5), 2031–6.

27. Suttapitugsakul, S.; Xiao, H.; Smeekens, J.; Wu, R., Evaluation and optimization of reduction and alkylation methods to maximize peptide identification with MS-based proteomics. Molecular BioSystems 2017, 13 (12), 2574–2582.

28. Müller, T.; Winter, D., Systematic Evaluation of Protein Reduction and Alkylation Reveals Massive Unspecific Side Effects by Iodine-containing Reagents. Molecular & Cellular Proteomics: MCP 2017, 16 (7), 1173–1187.

29. Kuznetsova, K. G.; Levitsky, L. I.; Pyatnitskiy, M. A.; Ilina, I. Y.; Bubis, J. A.; Solovyeva, E. M.; Zgoda, V. G.; Gorshkov, M. V.; Moshkovskii, S. A., Cysteine alkylation methods in shotgun proteomics and their possible effects on methionine residues. Journal of Proteomics 2021, 231, 104022.

30. Kuznetsova, K. G.; Solovyeva, E. M.; Kuzikov, A. V.; Gorshkov, M. V.; Moshkovskii, S. A., Modification of Cysteine Residues for Mass Spectrometry-Based Proteomic Analysis: Facts and Artifacts. *Biochemistry (Moscow)*, Supplement Series B: Biomedical Chemistry 2020, 14 (3), 204–215.

31. Saunders, C. C.; Stites, W. E., An electrophoretic mobility shift assay for methionine sulfoxide in proteins. Analytical Biochemistry 2012, 421 (2), 767–769.

32. Gundlach, H. G.; Moore, S.; Stein, W. H., The Reaction of Iodoacetate with Methionine. Journal of Biological Chemistry 1959, 234 (7), 1761–1764.

33. Vithayathil, P. J.; Richards, F. M., The Reaction of Iodoacetate with Ribonuclease-S. Journal of Biological Chemistry 1961, 236 (5), 1386–1389.

34. Lawson, W. B.; Gross, E.; Foltz, C. M.; Witkop, B., Alkylation and Cleavage of Methionine Peptides. Journal of the American Chemical Society 1962, 84 (9), 1715–1718.

35. Neumann, N. P., [56] Analysis for methionine sulfoxides. In Methods in Enzymology, Academic Press: 1967; Vol. 11, pp 487–490.

36. Lin, S.; Yang, X.; Jia, S.; Weeks, A. M.; Hornsby, M.; Lee, P. S.; Nichiporuk, R. V.; Iavarone, A. T.; Wells, J. A.; Toste, F. D.; Chang, C. J., Redox-based reagents for chemoselective methionine bioconjugation. Science 2017, 355 (6325), 597–602.

37. Kramer, J. R.; Deming, T. J., Reversible chemoselective tagging and functionalization of methionine containing peptides. Chem Commun (Camb) 2013, 49 (45), 5144–6.

38. van der Reest, J.; Lilla, S.; Zheng, L.; Zanivan, S.; Gottlieb, E., Proteome-wide analysis of cysteine oxidation reveals metabolic sensitivity to redox stress. Nat Commun 2018, 9 (1), 1581.

